# Adipocytes reprogram glucose metabolism in cancer cells promoting metastasis

**DOI:** 10.1101/2022.10.17.512437

**Authors:** Abir Mukherjee, Divya Bezwada, Malu Zandbergen, Francesco Greco, Chun-Yi Chiang, Medine Tasdemir, Johannes Fahrmann, Dmitry Grapov, Michael R. La Frano, Hieu S Vu, John W. Newman, Liam A. McDonnell, Luigi Nezi, Oliver Fiehn, Ralph J. DeBerardinis, Ernst Lengyel

## Abstract

In the tumor microenvironment, adipocytes function as an alternate fuel source for cancer cells. However, whether adipocytes influence macromolecular biosynthesis in cancer cells is unknown. Here, we systematically characterized the bi-directional interaction between primary human adipocytes and ovarian cancer (OvCa) cells using multi-platform metabolomics, imaging mass spectrometry, [^13^C]-glucose isotope tracing, and gene expression analysis. We report that omental tumor explants and OvCa cells co-cultured with adipocytes divert part of the glucose from glycolysis and TCA cycle towards glycerol-3-phosphate (G3P) synthesis. Normoxic HIF1α protein, stabilized by adipokines, regulate this altered flow of glucose-derived carbons in cancer cells, resulting in increased synthesis of glycerophospholipids (GPL) and triacylglycerols. Blocking adipocyte-induced HIF1α expression increases lipid peroxidation levels in cancer cells and sensitizes them to ferroptosis-mediated cell death. Subsequently, the knockdown of HIF1α or G3P acyltransferase 3 (a regulatory enzyme of GPL synthesis) reduced metastasis in xenograft models of OvCa. In summary, we show that in an adipose-rich tumor microenvironment, cancer cells generate G3P as a precursor for critical membrane and signaling components, thereby promoting metastasis. Targeting biosynthetic processes specific to adipose-rich tumor microenvironments might be an effective strategy against metastasis.

## INTRODUCTION

Several intra-abdominally metastasizing cancers (such as gastric, colorectal, and ovarian) have a propensity to seed to adipose tissues in the abdominal cavity, including the omentum, the small bowel mesentery, and the fat appendages on the large bowel (appendices epiploicae) ^1, 2^. The primary metastatic site for these cancers and the site for the largest tumor is the omentum, a large fat pad in the abdominal cavity overlying the small bowel. Colonization of the omental tissue marks an essential step in the metastatic cascade of ovarian cancer (OvCa), given that retrograde metastasis from the omentum is possible ^3^. The omentum, where adipocytes are the primary stromal cells, is distinct from the microenvironment of the ovary or fallopian tube where the OvCa originate. Therefore cancer cells have to rewire their metabolism to survive and proliferate in this lipid-rich environment.

We have shown that in epithelial OvCa, omental adipocytes contribute to the selective tropism of cancer cells through the secretion of adipokines ^4^. Upon interaction with the omental adipocytes, the OvCa cells transform them into “cancer-associated adipocytes” (CAA) ^1^, co-opting them for growth. To meet the energetic demands, OvCa cells promote lipolysis in adipocytes and take up lipids through the fatty acid receptor CD36, subsequently oxidizing these intracellular lipids ^5, 6^. However, it is unclear if the interaction of adipocytes with cancer cells results in additional metabolic changes in both the adipocytes and the cancer cells. Given, that increased biosynthesis of cellular components is an essential feature of proliferating cancer cells ^7^, the influence of adipocytes on such processes in cancer cells is yet to be determined.

To our knowledge, no systematic metabolic analysis of the bidirectional interactions between primary human adipocytes and cancer cells has been performed. Using untargeted global metabolomics and [^13^C]-glucose stable isotope tracing, we show that adipocytes redirect glucose utilization in cancer cells towards glycerol-3-phosphate (G3P), glycerophospholipid (GPL), and triacylglycerol (TAG) synthesis. The adipocytes induce “pseudo-hypoxia” through upregulation of HIFα in the cancer cells, thereby regulating the flow of glucose-derived carbons and promoting OvCa cell aggressiveness.

## RESULTS

### Multi-omics analyses show enhanced glucose utilization in both cancer and adipocyte compartments

We undertook an untargeted compartment resolved multi-omics approach to systematically study the metabolic interactions between adipocytes and OvCa cells. Adipocytes from non-cancerous human omentum (n=7) were isolated and co-cultured with OvCa cells (Fig. 1A). After (18hr) co-culture, media was collected, and each cell type was separated into cell pellets. The cell pellets and culture media were analyzed using three different metabolomics platforms to detect complex lipids, primary carbon metabolites, and oxylipins (Fig. 1B). Fold changes of the significantly altered metabolites in cancer and adipocyte compartments were grouped based on their biochemical and structural similarities, illustrated as network maps. Co-culture of cancer cells and adipocytes altered 134 metabolites in the cancer cells and 34 in the adipocytes (p < 0.05, mixed-effects ANOVA; Fig. 1C and Fig. S1A). Consistent with our previous finding that cancer cells use fatty acids from adipocytes for energy generation ^4^, we found increased triacylglycerol accumulation in cancer cells (n=33) (Fig. 1C) and decreased levels of fatty acids of various lengths in adipocytes (n=11) (Fig. S1A). Adipocytes also showed changes in pyruvate-derived metabolites (lactate, alanine) and TCA cycle intermediates (succinate and fumarate) (Fig. S1A) after co-culture with OvCa cells. A review of all metabolites in the co-cultured adipocytes suggests increased glucose oxidation *via* glycolysis and the TCA cycle. High glutamate levels in co-cultured adipocytes indicate simultaneous anaplerotic reactions ^8^, thereby maintaining the TCA cycle intermediate pool (Fig. S1A). The most striking change in the cancer cells upon co-culture was an increase in glycerophospholipids (n=76), with phosphatidylcholines (PC) being the most altered metabolite, followed by phosphatidylethanolamines (PE) (Fig. 1C). Glycerophospholipids are essential cell membrane components synthesized by incorporating fatty acids onto a glycerol backbone, with PC being the most abundant phospholipid ^9, 10^. However, glycerophospholipids can also enter the cell from the extracellular media/tumor microenvironment ^10^.

**Fig. 1.**
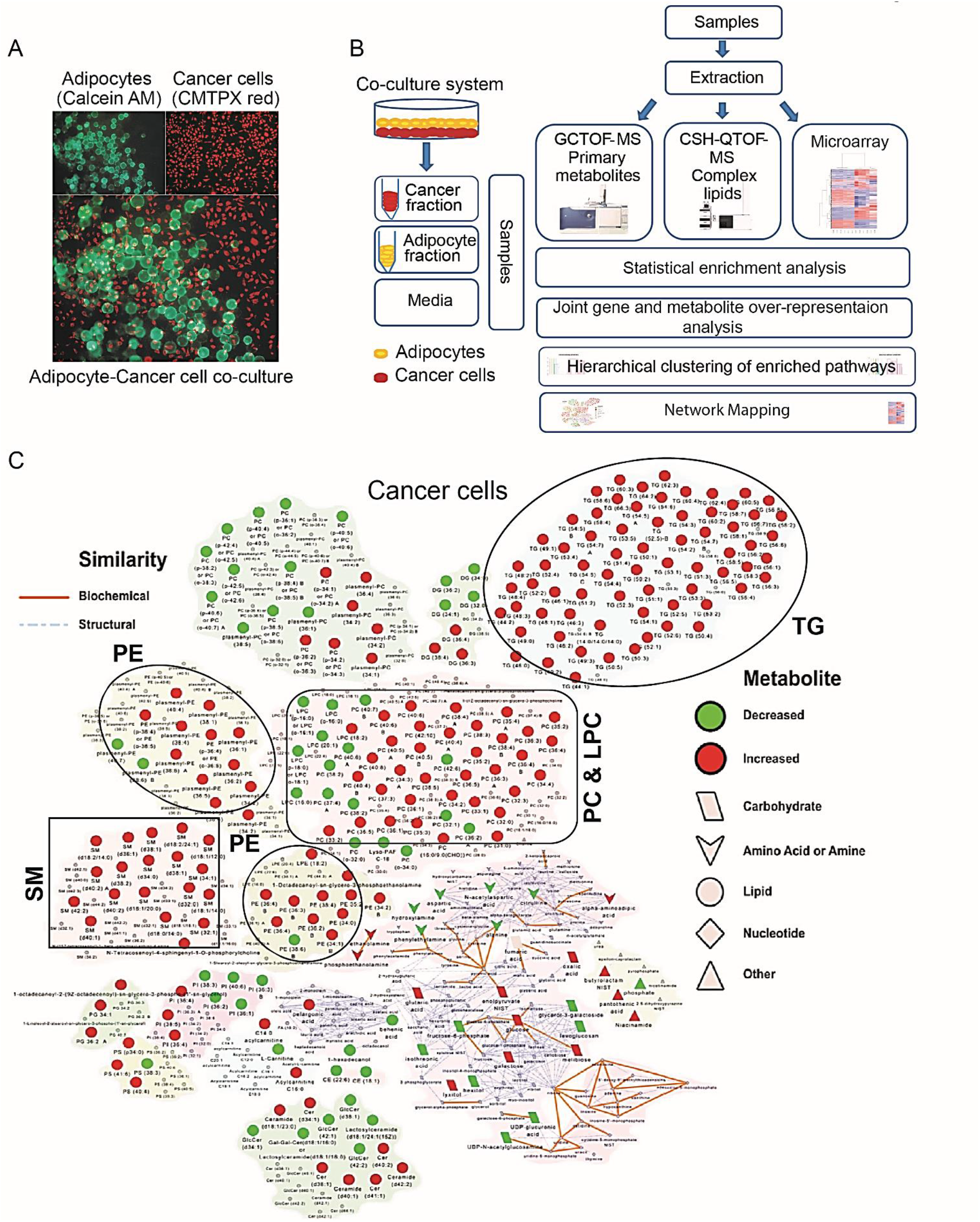
Adipocytes increase glucose metabolism and glycerophospholipids in ovarian cancer cells. **(A)** Co-culture of fluorescently (CMTPX) labeled ovarian cancer (OvCa) cells (red) with calcein AM-labeled primary human omental adipocytes (green). The adipocytes float on the surface because of their lipid content. **(B)** Schematic for multi-omics analyses of co-cultured adipocytes and OvCa cells. **(C)** Network map showing altered levels of metabolites (red = increase; green = decrease) in SKOV3ip1 cells with adipocyte co-culture. Biochemically and structurally similar metabolites are clustered. Orange lines connect metabolites in the same biochemical pathway, and blue lines connect structurally similar metabolites. Overall, glycerophospholipids (PC, phosphatidyl-choline; PE, phosphatidylethanolamine; SM, sphingomyelin; TG, triacylglycerol) are increased.

Our secretome analysis revealed that co-culture of adipocytes with OvCa cells altered metabolite levels in the media, adipocyte co-cultured cancer cells contributing more metabolites (n=83) than cancer cell co-cultured adipocytes (n=15; p < 0.05; Fig. S1B). Specifically, we observed increased monoacylglycerides, cyclooxygenase-derived metabolic products, and ceramides in the media from co-cultures (Fig. S1C). While some PCs detected in the media were adipocyte-derived, not all were taken up by cancer cells upon co-culture (Fig. S1D). PC (34:1), one of the most abundant phosphatidylcholines (PC) in cancer cells ^11^, is released in the media by adipocytes but its amount remained unchanged when co-cultured with cancer cells (Fig. S1D), suggesting that some of the PCs are synthesized within cancer cells.

Next, we explored the relationship between gene and metabolic networks of co-cultured adipocytes and cancer cells. There were 255 and 1065 differentially expressed genes in the adipocyte and cancer compartments, respectively (false discovery rate, FDR < 0.05; Fig. S2A, B, and Table 1). Gene set enrichment analysis (GSEA) revealed that co-culture leads to increased expression of cell cycle regulators in both cell types (Fig. S2C, D). In addition, a joint gene and metabolite over-representation analysis identified glycolysis as a critical pathway altered in both cellular compartments (Table 2) with increased gene expression of hexokinase 2 (HK2), glycerol-3-phosphate acyltransferase 3 (GPAT3), and lactate dehydrogenase (LDH) in cancer cells, and phosphofructokinase (PFK-p) in adipocytes (Fig. S2E, F). Therefore, in addition to lipid metabolism ^4, 5, 6^, co-culture with adipocytes enhances glucose metabolism in cancer cells.

### In omental metastasis, glucose-derived carbons are used for the synthesis of glycerol-3-phosphate and phospholipids

Our metabolomics data show that increased PC synthesis is the major change in adipocyte co-cultured cancer cells. The integrated gene expression/metabolite analysis also suggested increased glycolytic activity in cancer cells co-cultured with adipocytes (Fig. S2E). Therefore we explored if cancer cells use glucose to generate the glycerol backbone for phospholipids and triacylglycerols.

Consistent with our *in silico* analysis, adipocyte-derived conditioned media increased the extracellular acidification rate (ECAR) in two OvCa cell lines, showing significant increases in glycolytic rate (Fig. 2A, Fig. S3A, C). Additionally, this increase in glycolysis was accompanied by a decrease in oxygen consumption rate (OCR) (Fig. 2B, Fig. S3B, D), suggesting that glucose was incompletely oxidized. Subsequently, to determine how adipocytes influence glucose metabolism and to trace the flow of glucose-derived carbons in cancer cells, we performed stable isotope tracing analysis ^12^ on human omental explants obtained from three patients with advanced-stage high-grade serous OvCa. Cancer tissue adjacent to adipocytes was cultured *ex vivo* in media containing [U-^13^C]-glucose for 24hr, and the ^13^C enrichment profiles were mapped for central carbon pathway metabolites (Fig. 2C). We found high labeling in fructose-6-phosphate (m+6, range: 66-84%), lactate (m+3, range: 92-98 %), and reduced labeling in TCA cycle intermediates (range: 15-19% m+2 citrate and 10-11% m+2 succinate), which were consistent with our ECAR and OCR results, respectively (Fig. 2A, B). No significant labeling was observed in the other isotopologues of the TCA cycle intermediates (Fig. S3E). Together, these data suggest that under the influence of adipocytes, cancer cells rely on metabolites other than glucose to sustain the TCA cycle metabolite pool.

**Fig. 2.**
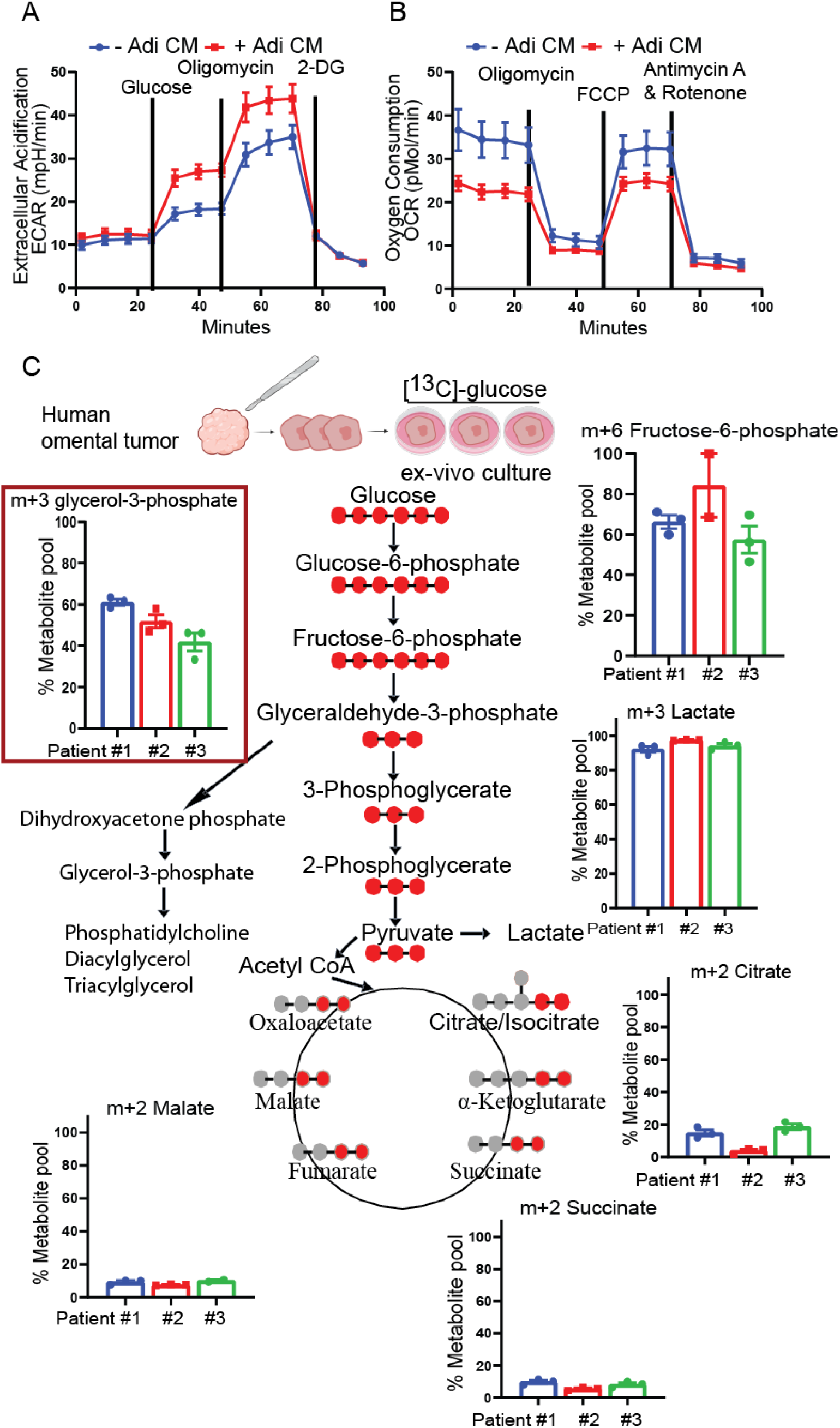
Adipocytes increase glycolysis but not oxidative phosphorylation in OvCa cells. SKOV3ip1 cells were treated with adipocyte conditioned media (Adi CM) for 18hr, followed by measurement of (**A**) extracellular acidification rate (ECAR), and (**B**) oxygen consumption rate (OCR) using Seahorse XF^e^96 analyzer. (n=3 independent experiments). **(C)**[^13^C]-glucose stable isotope tracing. Fresh omental tumor sections were incubated with [^13^C]-glucose for 24h, followed by mass-spectrometry to detect [^13^C]-carbon enrichment of the glycolytic pathway and TCA cycle.

Tumor explants also had high labeling (range: 42-61%) in glycerol-3-phosphate (G3P), the backbone for TAG and glycerophospholipids (Fig. 2C). Consistent with these results, we observed elevated m+3 labeling in lysophosphocholine 18:1 (Fig. S3F), with almost no labeling in the other isotopologues (m+4 to m+26) since fatty acids most likely contribute these carbons and are not derived from glucose. Overall, these findings indicate that in cancer tissue adjacent to adipocytes, glucose is partially utilized for the biosynthesis of the G3P.

### Glycerol-3-phosphate acyltransferase 3 regulates adipocyte-induced glycerophospholipid synthesis

Next, we explored the mechanism by which adipocytes increase glycerphospholipids in OvCa cells. Adipocyte co-culture increased glycerol-3-phosphate acyltransferase 3 (GPAT3) mRNA expression in cancer cells (Fig. S2E). GPAT3 is an enzyme upstream of glycerophospholipid and TAG synthesis that converts G3P into lysophosphatidic acid ^13^ (Fig. 3A). Consistent with GPAT3 regulation by adipocyte–cancer cell co-culture *in vitro*, we found increased GPAT3 protein expression in human omental metastasis. GPAT3 expression was detected at the invasive front of ovarian cancer cells pushing into human omental adipocytes (Fig. 3B), with lower expression in stroma and adipocytes. Using LC-MS to perform lipidomics, GPAT3 knockdown OvCa cells co-cultured with adipocytes resulted in reduced glycerophospholipids and TAGs, suggesting that adipocytes promote GPAT3-dependent GPL synthesis (Fig. 3C). Moreover, loss of GPAT3 reduced omental metastatic tumor burden *in vivo* (Fig. 3D), indicating that GPL synthesis in cancer cells is required for efficient omental colonization.

**Fig. 3.**
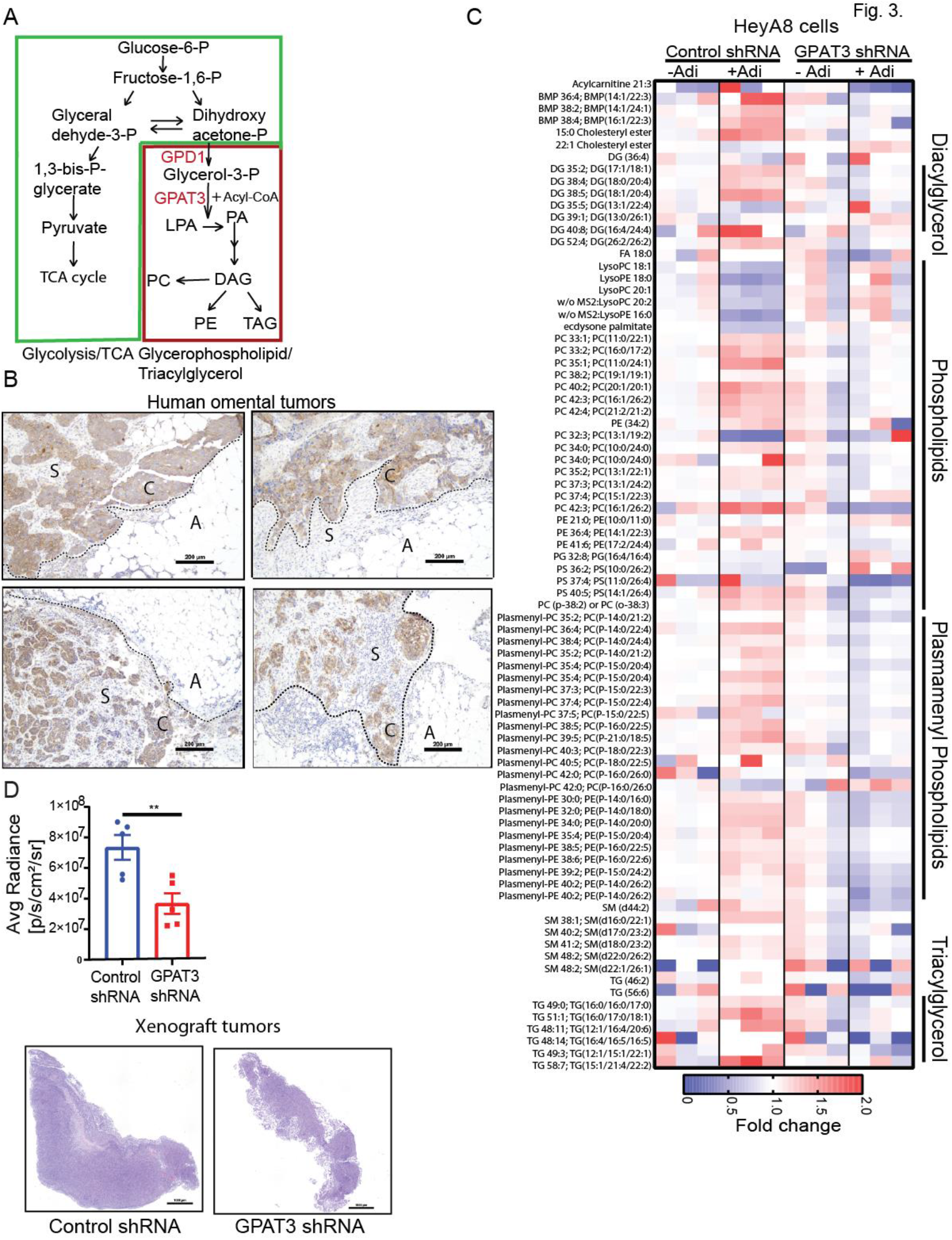
GPAT3 regulates glycerophospholipid synthesis. **(A)** Schematic. GPAT3 catalyzes the initial step in *de novo* glycerophospholipid and triacylglycerol synthesis. **(B)** GPAT3 expression in serial sections of patients with high-grade serous OvCa metastatic omental tumor. GPAT3 expression is more abundant in the epithelial tumor compartment. A – adipocytes, C-cancer, S-stroma, and dotted lines depict invasive front. (**C**) Lipidomics on control and GPAT3 shRNA OvCa cells co-cultured with omental adipocytes for 18hr. The heat map shows fold change of significantly altered lipid species (two-tailed t-test p < 0.05). Lipid species: DG Diacylglycerol, PC Phosphatidylcholine, Plasmenyl-PC Plasmenyl-Phosphatidylcholine, Plasmenyl-PE Plasmenyl-Phosphatidylcholine, TG Triacylglycerol, SM Sphingomyelin, BMP Bis(monoacylglycerol)phosphate, Lyso-PC Lysophosphatidylcholine, Lyso-PE Lysophosphatidylethanoloamine. (**D, top)** HeyA8-luciferase OvCa cells with stable control or GPAT3 knockdown were injected intraperitoneally into nude mice (n=5) and subsequently imaged using IVIS. Luciferase signal was quantitated to determine tumor burden (two-tailed t-test, ** p < 0.01, mean +/− SEM). (**D, bottom)** H&E stained sections of entire omental tumors

### Adipocyte-induced HIF1α shunts glucose-derived carbons to glycerophospholipid synthesis

Since glucose-derived carbons are used for G3P synthesis, we next sought to understand how adipocytes regulate this altered use of glucose in OvCa cells. Increased ECAR and reduced OCR (Fig. 2A, B) is an energetic profile frequently associated with hypoxia-inducible factor (HIF)1α expression ^14, 15^. Therefore, we considered that adipocytes possibly reprogram OvCa cell metabolism through HIF1α ^16^. Adipocyte co-culture increased HIF1α protein expression in multiple OvCa cell lines under normoxic conditions (Fig. 4A, Fig. S4A). HIF1α in the presence of oxygen undergoes prolyl hydroxylation and undergoes proteasomal degradation. Therefore, cells under normoxic conditions have a low abundance of the HIF1α protein. To determine whether adipocytes help stabilize HIF1α protein, we used a surrogate luciferase reporter plasmid, where the luciferase expression/activity is regulated by the oxygen-dependent degradation domain (ODD) of HIF1α. Therefore conditions/factors that stabilize HIF1α protein increase ODD-luciferase activity ^17, 18^. Stable transfection of the ODD-luciferase plasmid, followed by Adi CM treatment increased luciferase activity, suggesting that adipocytes stabilize HIF1α (Fig. 4B).

**Fig. 4.**
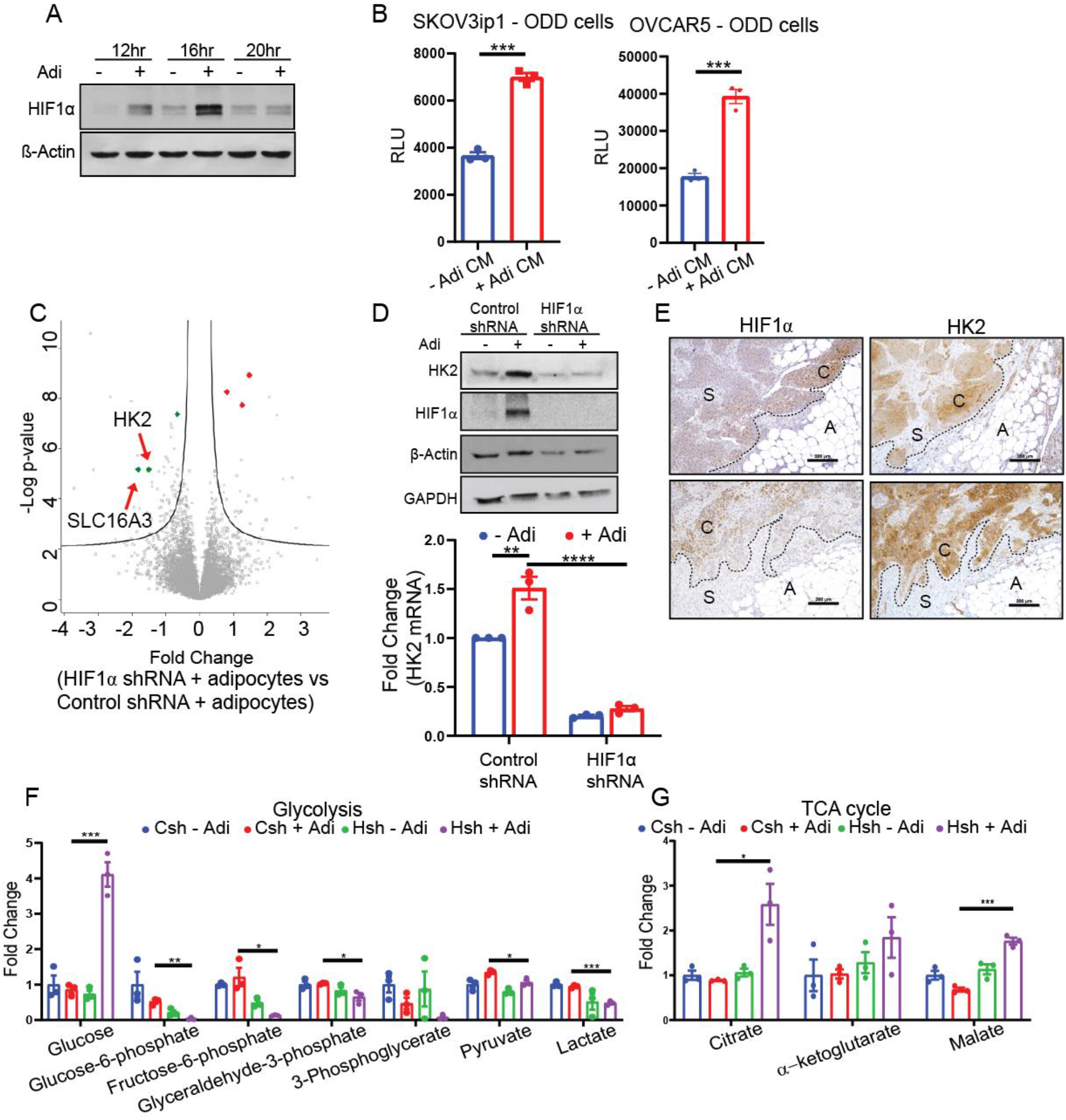
Adipocyte-mediated HIF1α expression regulates glucose utilization in cancer cells. **(A)** Immunoblot of HIF1α in SKOV3ip1 cells co-cultured with primary human adipocytes (Adi) for the indicated times. **(B)** SKOV3ip1 and OVCAR5 stably expressing the oxygen-dependent degradation domain (ODD) of HIF1α were treated with Adi CM for 6hr, and luciferase activity was determined using a luminometer (*** p < 0.001, two-tailed T-test). **(C)** Mass spectrometry of proteins extracted from OvCa cells stably transfected with shHIF1α and cultured +/− adipocytes. Volcano plot showing proteins regulated by adipocyte-induced HIF1α expression analyzed using a two-sided t-test with an FDR of 0.05 and an S0 value of 0.1). **(D)** Immunoblot (top) and qPCR (bottom) of HIF1α and HK2 in SKOV3ip1 HIF1α knockdown cells after adipocyte co-culture for 16hr. **(E)** Immunohistochemistry for HIF1α and HK2 using serial sections of human omental tumors from high-grade serous ovarian cancer patients. A – adipocytes, C-cancer, S-stroma, and dotted lines depict invasive front. (**F-G**) Metabolomics. Control and HIF1α knockdown SKOV3ip1 cells co-cultured with adipocytes for 18hr. Bar graphs show fold changes (compared to Control shRNA – Adi) in metabolites of **(F)** the glycolytic pathway and **(G)** the TCA cycle. Csh – Control shRNA, Hsh – HIF1α shRNA.

To identify the downstream targets of adipocyte-induced HIF1α in cancer cells, we performed a mass spectrometry-based proteomic analysis ^19^. HIF1α and scrambled short hairpin RNA (shRNA) were transduced into SKOV3ip1 OvCa cells (Fig. S4B) and co-cultured with primary human adipocytes. Proteomic analysis showed that HIF1α knockdown altered the adipocyte-induced proteome of OvCa cells, reducing 82 and increasing 22 proteins (FDR < 0.05 and S0 = 0.1; Fig. S4C, Table 3). Specific to glucose metabolism, HIF1α knockdown reduced protein expression of Hexokinase 2 (HK2) and the lactate/pyruvate transporter SLC16A3 (also known as MCT4) (Fig. 4C). Consistent with the proteomic data, adipocyte co-culture induced HK2 mRNA and protein levels in OvCa cells, dependent on HIF1α expression (Fig. 4D). Consistent with these results, we found increased HIF1α and HK2 expression in the epithelial tumor compartment adjacent to adipocytes in patients with omental metastasis (Fig. 4E). In addition, adipocyte-induced HIF1α also increased glycerol-3-phosphate dehydrogenase (*GPD1)* (Fig. S4D), an enzyme that converts the glycolytic intermediate dihydroxyacetone phosphate (DHAP) to glycerol-3-phosphate (Fig. 3A), thereby regulating the entry of glucose-derived carbons into glycerol-3-phosphate synthesis.

We next studied the effect of adipocyte-induced HIF1α on the glycolytic rate of OvCa cells. HIF1α knockdown reversed the effect of adipocytes on glycolysis (ECAR; Fig. S4E) and oxidative phosphorylation (OCR; Fig. S4F), thereby reversing the energetic profile we observed when co-culturing adipocytes with cancer cells (Fig. 2A, 2B). Further, HIF1α knockdown reduced glycolytic intermediates (Fig. 4F) and increased TCA cycle metabolites (Fig. 4G) with adipocyte co-culture, suggesting induction of “pseudo-hypoxia”^20^ by adipocytes. In addition, metabolomic data from adipocytes co-cultured HIF1α knockdown cells showed accumulation of glucose and reduced glucose-6-phosphate, which supports the hypothesis that adipocyte-induced glycolysis is primarily regulated by HK2 (Fig. 4F). HIF1α knockdown also blocked a majority of adipocyte-induced triacylglycerols (24/28), phosphatidylethanolamines (19/26), and ether-linked phosphatidylcholines (5/6), as well as other membrane lipids such as sphingomyelins (11/12) and ceramides (8/14) (Fig. S5). Together, these data suggest that adipocytes induce HIF1α expression in OvCa cells, divert glucose utilization towards TAG and glycerophospholipid synthesis.

Since adipocyte co-culture and Adi CM regulates glucose metabolism in cancer cells in a HIF1α-dependent manner, we sought to identify adipocyte-secreted soluble factors that regulate HIF1α expression in cancer cells. Fractionation of adipocyte CM using a size-exclusion filter revealed that the HIF1α inducing activity is present in the non-metabolite fraction of adipocytes CM (Fig. S4G). This finding was consistent with the secretome and intra-cellular analyses, showing no change in known metabolic regulators of HIF1α (fumarate, succinate, Fig. S4H). We and others have demonstrated that omental adipocytes secrete several cytokines/adipokines ^21^ that promote metastasis to the omentum ^4^. Treatment of OvCa cells with IL-6, IL-8, and MCP-1 increased HIF1α expression under normoxic conditions, with the most substantial effect from IL-6 (Fig. S4I). Neutralizing antibodies against IL-6 and MCP-1 abrogated adipocyte-induced HIF1α induction, while blocking IL-8 had no effect (Fig. S4J). Blocking downstream IL-6 signaling, using either STAT3 or JAK kinase inhibitors, led to a reduction in adipocyte CM-induced HIF1α, while a MEK inhibitor did not affect adipocyte-induced HIF1α expression (Fig. S4K). In summary, adipocytes induce HIF1α in cancer cells through an IL-6, JAK/STAT driven signaling pathway.

### Blocking glucose utilization inhibits omental metastasis

Increased HIF1α expression is a feature of aggressive and metastatic cells ^22^. We used a xenograft mouse model to determine the impact of dysregulated HIF1α on OvCa metastasis and to understand the flow of glucose-derived carbon in these tumors. Mice injected intraperitoneally with stable HIF1α knockdown OvCa cells had much smaller omental metastases (Fig. 5A). The HIF1α knockdown tumors had no change in apoptosis levels nor were there any differences in proliferation compared to the control tumors (Fig. 5B), indicating an alternate effect of HIF1α inhibition on OvCa cells. Ex-*vivo* co-culture of HIF1α knockdown cells with human omental tissue did not affect the ability of OvCa cells to adhere or proliferate (Fig. S6A) suggesting that HIF1α knockdown also don’t affect the initial steps of omental colonization ^23^.

**Fig. 5.**
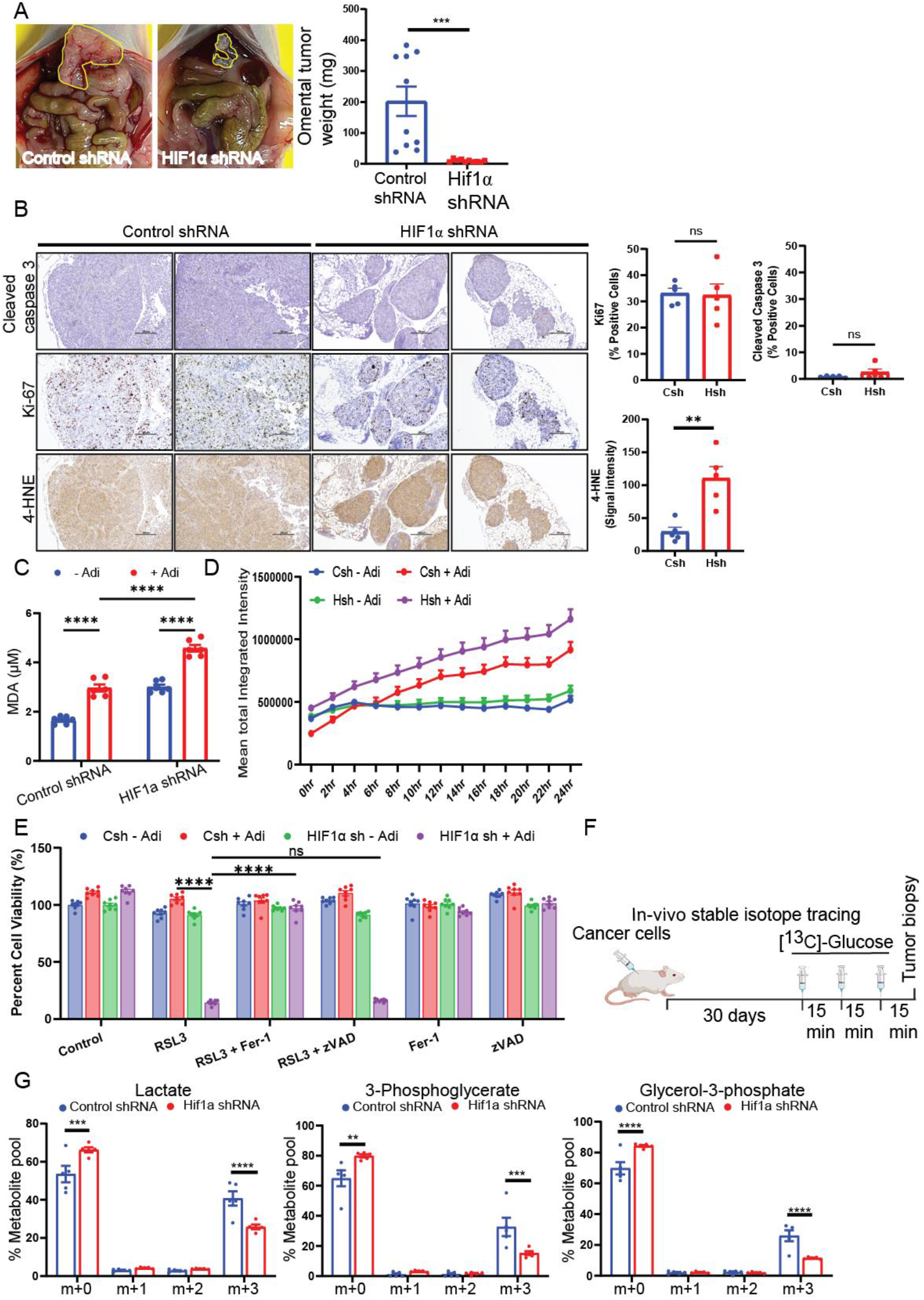
Synthesis of glucose-derived glycerol-3-phosphate is required for omental metastasis. Xenograft intraperitoneal metastasis model. **(A)** SKOV3ip1 HIF1α shRNA cells were injected intraperitoneally (n=10 mice per group), and tumor burden is shown (left panel) with sh control on the left and shHIF1α on the right with omental tumor outlined in yellow. Tumor burden was measured as omental metastatic weight (right panel). **(B)**Stable isotope tracing in tumor-bearing mice with HIF1α knockdown cancer cells. **(C)**[^13^C]-glucose derived labeling of glycerol-3-phosphate and glycolytic intermediates in omental tumors as in Figure 2C. **(D)** Cleaved caspase 3, Ki-67, and 4-HNE staining in serial sections of omental tumors generated from 5A (n=5 tumors per group were stained). Csh Control shRNA, Hsh HIF1α shRNA. (**E**) Lipid peroxidation: Control and HIF1α knockdown cells were co-cultured with adipocytes for 18hr and intra-cellular malondialdehyde (MDA) levels determined. (**F-G**) Control and HIF1α shRNA transduced SKOV3ip1 cells were treated with Adi CM. (**F**) Cells were stained using Bodipy 581/591 C11 dye and fluorescent emissions (at 520nm) were measured every couple of hour using IncuCyte to quantify lipid ROS production. (**G**) MTT assay carried out to determine the viability of cancer cells after treatment with the indicated small molecule compounds.

Previously we showed that adipocyte co-culture increased lipid peroxidation in OvCa cells ^5, 6^. Using 4-hydroxynonenal (4-HNE) adducts as a read-out for lipid peroxidation, we found increased staining intensities in HIF1α knockdown tumors (Fig. 5B). HIF1α knockdown cells cancer cells co-cultured with adipocytes showed significantly higher amount of malondialdehyde (MDA), a marker of lipid (polyunsaturated fatty acid) peroxidation (Fig. 5C). Both *in-vivo* and *in-vitro* data show that blocking HIF1α expression in cancer cells in the presence of adipocytes increased lipid peroxidation in cancer cells. Since uncontrolled lipid peroxidation causes ferroptosis ^24^ we next explored if knockdown of HIF1α sensitized OvCa cells towards ferroptosis mediated cell death. While we observed an increase in lipid ROS levels in control shRNA cells with Adi CM, the knockdown of HIF1α exacerbated the levels (Fig. 5D, and S6B). GPX4 is the primary enzyme responsible for removing membrane lipid peroxides ^25^ therefore we measured the survival capacity of OvCa treated with RSL3 (a GPX4 inhibitor) ^26^. We found that in the presence of Adi CM, HIF1α knockdown cells were specifically sensitive to the inhibition of GPX4 (Fig. 5E). Further, reversal of cell viability by ferrostatin (a ferroptosis inhibitor ^26^) but not by zVAD-FMK (a pan-caspase inhibitor) showed that HIFα knockdown cells undergo ferroptosis, an apoptosis-independent cell death (Fig. 5E).

To evaluate the effect of HIF1α knockdown on glucose utilization *in vivo*, tumor-bearing mice (control/scrambled or HIF1α shRNA) were infused with [U-^13^C]-glucose ^27^. Upon necropsy, omental tumors were excised, and carbon distribution was measured using mass spectrometry (Fig. 5F). HIF1α knockdown reduced the flow of glucose into glycolysis (lactate m+3, and 3-phosphoglycerate m+3) and significantly reduced m+3 labeling of glycerol-3-phosphate (Fig. 5G). These data provide evidence for regulation of glycerol-3-phosphate synthesis by HIF1α *in-vivo*.

### Spatial distribution of glycerophospholipids in human omental metastatic samples

To clinically validate the effect of adipocytes on glycerophospholipids synthesis, we carried out imaging mass spectrometry (IMS) analysis on sections of human omental tumors. Based on our previous studies we observed that the effect of adipocytes on cancer cells is dependent on their proximity to adipocytes. Therefore, to determine the adipocyte-induced phospholipid changes in cancer cells, we evaluated the differences in phospholipid species in tumor cells at the leading edge invading into adipocytes compared to cancer cells farther away (more than 200μm from the adipocytes). Fresh frozen omental tumor sections from three high-grade serous ovarian cancer (HGSOC) patients were coated with norharmane, a dual polarity matrix ideal for phospholipid analysis ^28^. Subsequently, images of metabolites (ions ranging between 150-2000 m/z) were taken using MALDI (Fig. 6A), and metabolites in cancer cells adjacent to the adipocytes were compared to cancer cells further away (Fig. 6B and 6C). To identify metabolites that distinguished the two groups of cancer cells, we carried out a receiver operating curve (ROC) analysis (Table 4). The analysis showed that cancer cells adjacent to adipocytes had a higher abundance of PCs, while those at a distance had more phosphatidylinositol (PI) (Fig. 6D and 6E). This corroborates our observation using primary adipocytes that adipocytes adjacent to cancer cells increase PCs in tumor cells.

**Fig. 6.**
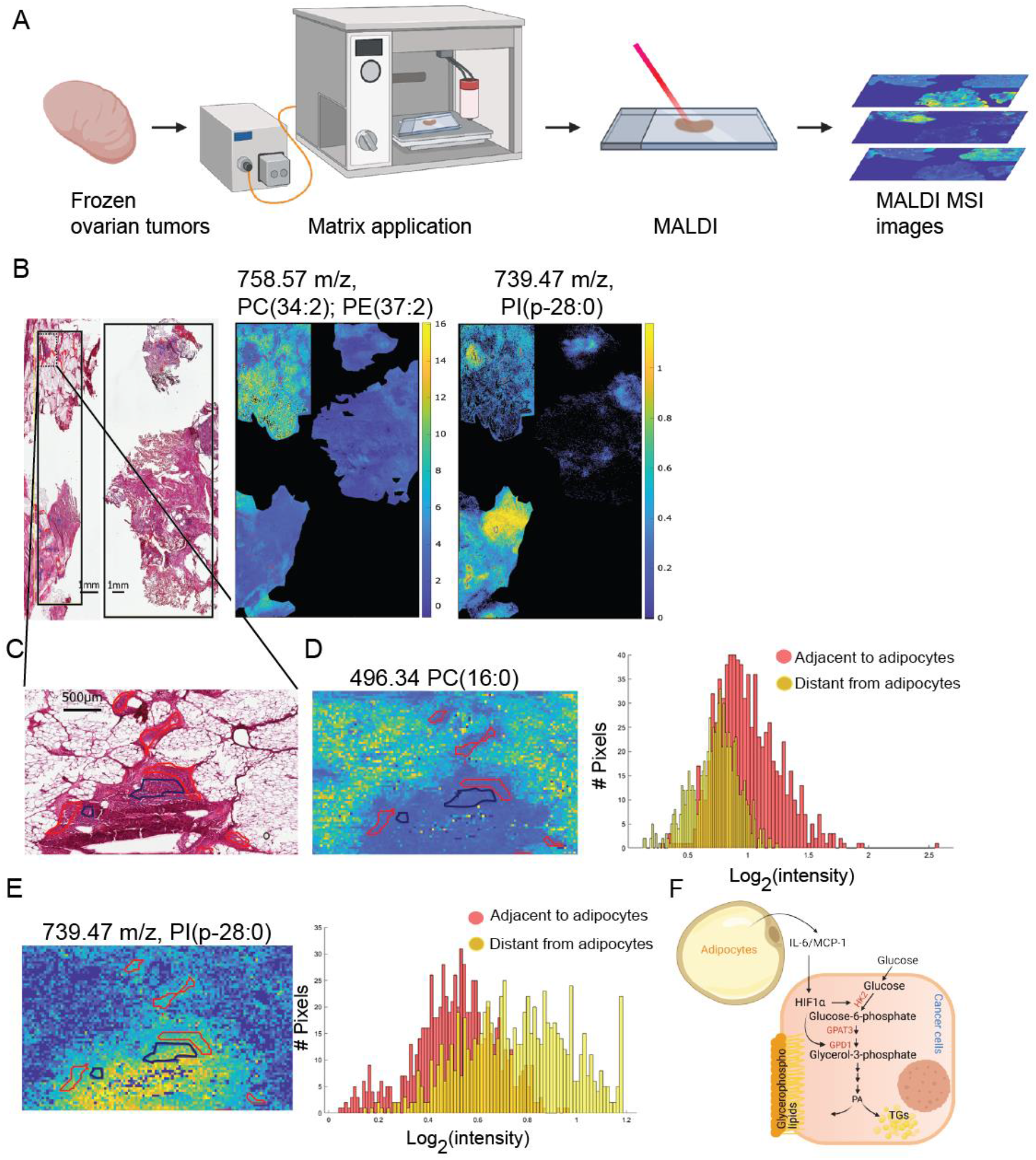
Cancer cells adjacent to adipocytes have increased phosphatidylcholine. Spatial analysis of glycerophospholipids using imaging mass spectrometry (IMS). **(A)** Schematic for IMS analysis of ovarian high grade serous tumor samples. **(B-C)**, Histology and MALDI IMS analysis of frozen omental tumor sections. **(B, top panel)** Hematoxylin and Eosin stained tissue with annotated tumor regions (black box) that were analyzed using MALDI. **(B, bottom panel)** Spectral distribution of ions (496.34 m/z and 739.47 m/z) corresponding to the selected area (black box) on the H&E slide. **(C)** Enlarged section of the tumor area (used for IMS analysis), showing two groups of cancer cells, adjacent (red) or distant (blue) from adipocytes. **(D-E)** Optical images using MALDI IMS of phosphatidylcholine PC (16:0) or Phosphatidylethanolamine (PE) 37:2 (higher adjacent to adipocytes as measured by the AUC-orange), (**D)** and Phosphatidylinositol (PI) p-28:0 (lower adjacent to adipocytes as measured by the AUC-orange) **(E)** Corresponding to the H&E section shown in panel **(C)**. MALDI images of ions are shown at the top of the panel, and the pixel intensities of the ions in the selected area are at the bottom. (n=3 human omental tumor samples were analyzed). **(F)** Illustration of the adipocyte-ovarian cancer cell interactions leading to altered glucose utilization in OvCa cells. Adipocytes stabilize HIF1α protein in OvCa cells. Subsequently, HIF1α transactivates hexokinase 2 (HK2) and glycerol-3-phosphate dehydrogenase (GPD1). The combined activity of these two enzymes increases glycerol-3-phosphate levels in OvCa cells, forming the backbone for the synthesis of triacylglycerol (TGs) and glycerophospholipids.

## DISCUSSION

We found that adipocyte co-culture activates HIF1α signaling in OvCa cells, shunting glucose towards glycerol-3-phosphate and phospholipids synthesis, thereby supporting metastatic colonization (Fig. 6F). The symbiotic relationship between adipocytes and cancer cells explains why omental tumors are the most common and the largest, in abdominally metastasizing cancers such as ovarian and gastric ^29^. Tumor cells interacting with adipocytes in the microenvironment induce lipolysis in adipocytes, and fatty acids transferred to cancer cells are used for energy production ^1^. In addition, surgical removal of the omentum at early or metastatic disease reduces the recurrence risk in abdominally metastasizing tumors ^30, 31^. In these tumors, interaction of omental adipocytes and cancer cells influence disease outcomes. Our systematic analyses of the metabolic changes that occur upon the interaction of adipocytes with cancer cells show that adipocytes also activate glycolysis in cancer cells and shunt glucose towards glycerol-3-phosphate and glycerophospholipid synthesis (Fig. 2C). Glycerophospholipids, regulated by GPAT enzymes, are essential components of cellular lipid membranes and signaling molecules that are important for cell division ^13^. Therefore, not surprisingly, increased glycerophospholipids levels (PCs and PEs) are observed in bladder, kidney, and colorectal cancers ^32, 33^.

We show that omental adipocytes stabilize HIF1α via IL-6 and MCP-1 independent of hypoxia, which diverts glucose-derived carbons to glycerol-3-phosphate, thereby generating the backbone for glycerophospholipids and TAG synthesis. In our experimental approach, under adipocyte-induced ‘pseudo-hypoxia’, cancer cells are not reliant on glucose for fatty acid synthesis (Fig. S3F). There is no need for the metastatic cancer cells to synthesize fatty acids, given the abundance of lipids in the adipocyte-rich tumor microenvironment. In adipocyte-rich tumor microenvironments cancer cells have a distinct glucose metabolic profile. This contrasts to hypoxic conditions, where glucose generates both glycerol-3-phosphate backbone and saturated fatty acids for phosphatidylcholine synthesis ^11^.

Several mechanisms contribute to the reduced tumor burden observed in HIF1α knockdown xenografts. First, HIF1α knockdown xenografts showed a reduced flow of glucose into glycerol-3-phosphate. This is critical for TAG and glycerophospholipid synthesis as building blocks for new membranes in dividing cancer cells and for reducing the adverse effects of toxic free fatty acids ^34, 35, 36^. Secondly, HIF1α knockdown xenograft tumors showed increased lipid peroxidation, known to be toxic to tumor cells ^37^. Thirdly, adipocyte co-cultured HIF1α knockdown cells have increased lipid ROS and lipid peroxidation productions, making them uniquely susceptible to GPX4 inhibition. Consistent with our findings, in fibrosarcoma cells, reduced HIF1α expression increases lipid peroxidation and subsequently ferroptosis ^38^. While it is yet to be determined if HIFα knockdown reduces the capacity of OvCa cells to remove toxic membrane-derived lipid peroxides, we observed accumulation of polyunsaturated plasmalogens such as PE P-40:6 (known to induce ferroptosis ^39^) in HIF1α knockdown cells co-cultured with adipocytes.

This study expands our current understanding of cancer-associated adipocytes by demonstrating a complex symbiotic relationship with cancer cells and a role in completely reprogramming cancer cell metabolism ^2^. In addition to supplying fatty acids to cancer cells for β-oxidation, adipocytes help reprogram signaling pathways through HIF and provide building blocks for membranes and lipid mediators that allow cancer cells to divide and proliferate. Adipocytes support the extreme energy requirements of cancer cells through a combination of adipokine secretion, lipolysis, and, as we show in this study, the reprogramming of glucose metabolism. Consequently, interfering with the adipocyte-cancer cell interactions could slow metastasis and, if implemented early after the initial tumor debulking surgery, might delay the frequently seen recurrence of intra-abdominally recurring tumors to adipose tissue.

## MATERIALS AND METHODS

### Cell lines and reagents

SKOV3ip1, HeyA8, and OVCAR5 cells were cultured as described previously ^5^. TYKnu cells were cultured in eagle’s minimal essential media (MEM) supplemented with 10% fetal bovine serum. All cells were cultured in an incubator maintained at 37°C with 5% CO_2_. Cell lines were regularly tested for mycoplasma and genotyped for authentication purposes (IDEXX Bioresearch short tandem repeat marker profiling). Neutralizing antibodies against human IL-6 (cat# MAB206), IL-8 (cat#MAB208), MCP-1 (cat#AB279) and isotype specific control were purchased from R&D systems and used at a concentration for 100μg/ml. Recombinant human cytokine/chemokines IL-6 (cat# 200-06), MCP-1 (cat# 300-04) and IL-8 (cat# 200-08) was purchased from PeproTech and used at a concentration of 10ng/ml. 1S, 3R-RSL3 (RSL3) was purchased from Sigma-Aldrich (cat#SML2234) and zVAD-FMK (zVAD) from Selleckchem (cat#S7023). Poly-L-lysine solution 0.1%, norharmane, HPLC grade chloroform, and LC-MS grade methanol were purchased from Sigma-Aldrich (Saint Louis, MO, USA). Histology-grade solvents and Eosin G or Y (alcoholic solution 0.5%) were purchased from DiaPath S.p.A. (Martinengo, Italy); Harris hematoxylin was obtained from VWR International, LLC (Radnor, PA, USA). LC-MS grade water was purchased from Thermo Fisher Scientific (Rockford, IL, USA), and indium-tin oxide (ITO) coated glass slides for MALDI MSI were obtained from Bruker Daltonics (Bremen, Germany).

### Isolation of primary adipocytes and cell culture

Human primary omental adipocytes were isolated as previously described ^4, 40^. Briefly, freshly excised omental tissues were digested using 0.2% collagenase type I enzyme for 1hr and separated from the stromal vascular fraction using 250 μm nylon mesh. Adipocytes were repeatedly washed with media (DMEM: F12 media with 1% streptomycin and 1% penicillin) and centrifuged to remove contaminating cells. Isolated adipocytes were cultured in DMEM: F12 media supplemented with 0.1% fatty acid free BSA (Roche, NY), 100 U/ml penicillin, and 100 μg/ml streptomycin. Co-culture of adipocytes with cancer cells was carried out in 1:5 packed cell adipocyte volume (PCV) to serum-free media (SFM). To generate adipocyte-conditioned media (CM), adipocytes were cultured in DMEM: F12 media for 72hr, after which media was removed and filtered through a 0.8 μm syringe filter. Size-based fractionation of Adi CM was carried out by using a 3 kd filter (Amicon, Millipore) and centrifuged at 4000 rpm for 45min at 4°C. Isolated adipocytes and SKOV3ip1 cells were stained with Calcein AM (10 μM) in DMEM: F12 media for 20min, washed with PBS once, and images were taken using Zeiss Axio Observer.A1 microscope and images processed using Axios Vision LE software.

### Human tissue

All patients have consented before surgery, and patient-derived tissues were obtained fresh or were paraffin-embedded according to approved Institutional Review Board (IRB) protocol. High-grade serous ovarian cancer tumor samples were obtained from patients with clinical stage III and IV disease (FIGO) undergoing primary debulking surgery at the University of Chicago hospital. For adipocyte co-culture experiments, human omental tissues were obtained from patients with benign, non-cancer related conditions (e.g. incontinence, fibroids).

### Mouse studies

Intraperitoneal xenograft experiments were performed by injecting SKOV3i1p1 (5 million cells/mouse) or HeyA8-luciferase (1 million cells/mouse) cells into female immunocompromised athymic nude mice (Envigo), and tumor burden determined after 4 and 2 weeks post injection, respectively^5^. Tumor burden in mice injected with GPAT3 knockdown cells was assessed by injecting 100μl D-luciferin (30 mg/ml) 10 minutes before imaging using IVIS spectrum *in vivo* imaging system. For the *in vivo* [^13^C]-glucose tracing experiment, 5 million stable SKOV3ip1 cells (transduced with either control shRNA or shRNA targeting HIF1α) were injected into female athymic nude mice (Envigo), and tumors were allowed to establish for 4 weeks. Mice were injected three times with 1.34 mM [^13^C]-glucose solution in PBS (80 μl) at 15 minute intervals. Omental tumors were excised and snap frozen in liquid nitrogen. All animal studies were approved by the Institutional Animal Care and Use Committee at the University of Chicago.

### [^13^C]-Glucose tracing analysis

Glucose enrichment analysis on human omental explant tissues (from three HGSOC patient, three sections were analyzed from each tumor) and mouse omental tissues was carried out as described previously ^27, 41^. Omental tumor biopsies from HGSOC patients were sectioned (0.5-1 mm) and incubated in DMEM supplemented with 10 mM [U-^13^C]-glucose (Cambridge Isotope Laboratories) and 2 mM glutamine. Explants were incubated for 24hr in an incubator maintained at 37°C with 5% CO_2_, with gentle rocking. Metabolites from mouse omentum isolated from xenograft experiments (as described above) and *ex vivo* cultures of human omental tumors were extracted using 80% cold methanol. Protein estimation was carried out using a BCA assay (Pierce, Thermo Scientific), and equal protein amounts were dried using a speed-vac. Data acquisition was performed by reverse-phase chromatography on a 1290 UHPLC liquid chromatography (LC) system interfaced to a high-resolution mass spectrometry (HRMS) 6550 iFunnel Q-TOF mass spectrometer (MS) (Agilent Technologies, CA). The MS was operated in positive and negative (ESI+ and ESI-) modes. Analytes were separated on an Acquity UPLC® HSS T3 column (1.8 μm, 2.1 × 150 mm, Waters, MA). The column was kept at room temperature. Mobile phase A composition was 0.1% formic acid in water, and mobile phase B composition was 0.1% formic acid in 100% ACN. The LC gradient was 0 min: 1% B; 5 min: 5% B; 15 min: 99% B; 23 min: 99% B; 24 min: 1% B; 25 min: 1% B. The flow rate was 250 μL min^−1^. The sample injection volume was 5 μL. ESI source conditions were set as follows: dry gas temperature 225 °C and flow 18 L min^−1^, fragmentor voltage 175 V, sheath gas temperature 350 °C and flow 12 L min^−1^, nozzle voltage 500 V, and capillary voltage +3500 V in positive mode and −3500 V in negative. The instrument was set to acquire over the full *m/z* range of 40–1700 in both modes, with the MS acquisition rate of 1 spectrum s-1 in profile format.

Raw data files (.d) were processed using Profinder B.08.00 SP3 software (Agilent Technologies, CA) with an in-house database containing retention time and accurate mass information on 600 standards from Mass Spectrometry Metabolite Library (IROA Technologies, MA), which was created under the same analysis conditions. The in-house database matching parameters were: mass tolerance 10 ppm; retention time tolerance 0.5 min; co-elution coefficient 0.5. The peak integration result was manually curated in Profinder for improved consistency and exported as a spreadsheet (.csv).

### Metabolomic analysis

Primary omental adipocytes isolated from 7 consenting patients were co-cultured with SKOV3ip1 cells for 18hr (1:5 PCV, adipocytes: FM), after which adipocytes, cancer cells, and media were collected for metabolomic analysis. Equal number of cells and an equal volume of media were aliquoted for analysis. Sample extraction and untargeted metabolomics analysis of primary metabolites and complex lipids were carried out as previously described ^42, 43^. Analysis of oxylipins, endocannabinoids, and ceramides (in the media) were constructed using GC and LC-MS platforms ^44, 45^ at the West Coast Metabolomics Center, University of California, Davis.

### Microarray analysis

SKOV3ip1 cells were co-cultured with primary omental adipocytes for 8hr. Cancer cells and adipocytes were separated and washed several times with PBS. Cells were lysed using Trizol (Life Technologies), and total RNA was isolated according to the manufacturer’s protocol using RNeasy kit (Qiagen), following in column DNase treatment. Microarray analysis was carried out using the Illumina BeadChipHT-12v4 expression array as previously described ^5^. Data processing was carried out using Illumina Genome studio v2011.1 and Partek Genomics suite v6.6. Average signal intensities were background subtracted, log_2_ transformed, and quantile normalized. Customized data processing scripts were developed in R version 3.3.0. Each experimental group had three replicates, and differentially expressed genes were identified as genes with a q-value of 0.05 and a fold change of 2 and above.

### MALDI mass spectrometry imaging

Omental tumor samples were obtained from three HGSOC patients. Samples were rinsed in physiological saline and immediately frozen and stored at −80°C until analysis. Tissue sections (12 μm) were cut using a Leica CM1950 cryostat (Leica, Wetzlar, Germany) and thaw mounted onto polylysine coated ITO conductive slides. Slides were stored at −80°C until analysis. The tissue sections were thawed under vacuum for 15 minutes before matrix application. Norharmane, a dual polarity matrix (7 mg/mL in CHCl_3_: MeOH 70:30) was applied in 8 layers (2 layers at 5 μL/min followed by 6 layers at 10 μL/min) using a SunCollect (SunChrom, Friedrichsdorf, Germany) matrix spraying system (4 bar, 50 mm z axis height). Tissue sections were dried under vacuum for 15 minutes before MALDI MSI data acquisition. MALDI MSI images were acquired using an EP-MALDI source^46^ (Spectroglyph, LLC., Kennewick, WA, USA) equipped with a 349 nm laser (Spectra-Physics, Santa Clara, CA, USA), coupled to an Orbitrap QExactive Plus (Thermo Scientific). The laser was set to 1.65 A and 500 Hz, the ion source pressure was 7.2 Torr; MSI pixel size was 35×35 μm. Mass spectra were acquired in the range of 150-2000 *m/z* at 70 000 resolving power. Position files were aligned to the raw file using Image Insight (v. 0.1.0.11550, Spectroglyph, LLC). We analyzed a total of 40 and 34 tissue regions (in the three tumor sections) comprising of cancer cells adjacent and distant to the adipocytes, respectively.

### Imaging mass spectrometry - Image preprocessing and data analysis

Raw spectra of the MSI dataset were converted into mzXML using RawConverter ^47^. A modified version of the script ORBIIMAGEmzXML2Tricks (v. 0.10, G. Eijkel) was used to produce a basepeak spectrum from each MSI dataset. Basepeak spectra of all the samples were summed to obtain a global basepeak spectrum. Peak picking was then performed on the global basepeak spectrum, and the resulting peak list was used to extract the MS images from each sample’s dataset using ORBIIMAGEmzXML2Tricks. MSI datacubes were imported into MATLAB R2019b (MathWorks, Natick, MA, USA) for image preprocessing and data analysis. Each MSI dataset was processed separately to remove peaks that negatively correlated with the tissue, each peak list was deisotoped, and the intensity of pixels outside the tissue was set to zero. A merged datacube was produced by selecting only those peaks that were detected in all tissue sections being compared and then using spatial offsets to place all datasets within the same coordinate space. The resulting datacube was TIC normalized, and bright spots with an intensity over the 99.9^th^ percentile were removed.

### Imaging mass spectrometry - Histological staining and annotation

Tissue sections were stained with hematoxylin and eosin (H&E) after MSI data acquisition. Briefly, residual MALDI matrix was removed with ethanol washes (2X 30s each) followed by rehydration and H&E staining. Optical images of the histologically stained tissues were recorded using an Aperio CS2 scanner at 40x magnification (Aperio Technologies Inc., Vista, CA, USA) and annotated using Aperio ImageScope (v 12.2.2.5015, Aperio Technologies Inc.). Ovarian cancer cells adjacent to adipocyte regions were defined as tumor cells closer than 200 μm to adipocytes; these tumor cells were shown to present intracellular lipid staining in a previous work^4^. Ovarian cancer cells further than 200 μm from adipocytes were annotated as distant from adipocytes. Histological images were imported in MATLAB and co-registered to the MALDI MSI dataset. Annotated regions were imported in MATLAB, converted into masks and aligned to the MSI dataset, in order to perform region-based analysis^48^. Region masks were merged together using the same spatial offsets of the MSI dataset to obtain a global mask that can be applied to all MSI images at once.

### MALDI MSI data analysis

ROC curve analysis was used to identify *m/z* that differentiate ovarian cancer cells adjacent to adipocytes from those distant from adipocytes. ROC curve analysis was performed by comparing the log2 transformed intensities of the pixels. Only *m/z* features with a non-zero value in at least 30% of the pixels of the regions were considered for the analysis. To take into account any bias that may arise from the different dimensions of the regions, ROC analysis was performed on randomly selected pixels from the regions, using the number of pixels of the smallest area. ROC analysis was performed in MATLAB using the function *perfcurve;* ions corresponding to an area under the curve (AUC) greater than 0.7 (increased intensity in ovarian cancer cells adjacent to adipocytes) and lower than 0.3 (increased in ovarian cancer cells distant from adipocytes) were selected. Selected *m/z* were assigned based on the mass accuracy using METLIN^49^ (Δ=10 ppm, [M+H]^+^,[M+Na]^+^,[M+H-H_2_O]^+^ adducts). Lipids are presented using the lipid species level notation ^5, 50^.

### Quantitative real-time RT-PCR analysis

To determine the changes in mRNA expression, cancer cells were plated in 6-well plates and allowed to attach for 36hr after which the media was removed, cells washed with PBS (twice), and co-cultured with primary adipocytes (1:5 PCV ratio) for 12hr^6^. Adipocytes were removed by pipetting, and cancer cells were washed with PBS (3-5 times) to remove the remaining adipocytes. Cells were then lyzed using Trizol (Life Technologies) and total RNA extracted (per manufacturer’s protocol). cDNA synthesis was carried out using 1 μg total RNA using high-capacity cDNA synthesis kit (Applied Biosystems). Subsequently, qPCR was carried out using the 7500 Real-Time PCR System (Applied Biosystems) with probes for HK2 (Hs00606086_m1), GPAT3 (Hs00262010_m1), GPD1 (Hs01100039_m1), and normalized to GAPDH (Hs02758991_g1) as a loading control. Relative changes were calculated using 2^−ΔΔCt^ method, and a p-value of < 0.05 was considered significant (t-test).

### Immunoblotting

Cell lysis was carried out using RIPA buffer supplemented with protease and phosphatase inhibitors. Subsequently, protein estimations were performed using BCA protein assay kit (Pierce, Thermo Scientific). Equal protein amounts were resolved on SDS gels (4-20%), transferred onto nitrocellulose membrane, and blotted for antibodies against specific proteins HIF1α (BD biosciences, cat# 610958, 1:1000), HK-2 (Cell Signaling Technology, cat# 2867, 1:1000), p-STAT3 Tyr 708 (Cell Signaling Technology, cat# 9131, 1:1000), total STAT3 (Cell Signaling Technology, cat# 9139, 1:1000), GAPDG (Cell Signaling Technology, cat# 5174, 1:1000) and β-Actin (Millipore Sigma cat# A5441, 1:3000).

### Immunohistochemistry

Immunohistochemistry was carried out as previously discussed ^4^. Serial sections of omental tumors from HGSOC patients were stained using antibodies against HIF1α (BD biosciences, cat# 610958, 1:50), HK2 (Cell Signaling Technology, cat# 2867, 1:50), GPAT3 (Novus cat# NBP-1-93629, 1:250), Ki-67 (Thermo Scientific Labvision, Cat# RM-9160, 1:400), cleaved caspase 3 (Cell Signaling Technology, cat# 9661, 1:200) and 4-HNE (Abcam cat# ab46545, 1:300).

### Lentivirus production and RNA interference

293T cells were transfected with packaging plasmids (8 μg of pCMV-dR8.2 dVPR and 1 μg pCMV-VSV-G, a gift from Robert Weinberg, Addgene plasmid #8455 and 8454, respectively) along with shRNA plasmids (8 μg) using Lipofectamine 2000 (Thermo Fisher Scientific) transfection reagent. Viral supernatants were collected 48hr after transfection and added to cell lines after filtration using 0.45 μm syringe filters. Cancer cell lines were transduced with lentivirus containing shRNA targeting HIF1α (TRCN0000003810) and GPAT3 (TRCN0000162335) to generate stable lines.

### Omental explant assay

Human omental tissue was collected and ex-vivo culture carried out ^5^ to determine the effect of HIF1α knockdown on the colonization capacity OvCa cells. 10-mm punch biopsies of omental tissue (3 per group) were added to a 24-well ultra-low attachment plate. One million scramble/control or HIF1a targeting shRNA transduced SKOV3ip1 cells were stained using CellTracker™ Red CMTPX Dye (Thermo Fisher Scientific) and added to each well. After 18 (adhesion) and 72 hours (proliferation) culture, the cells were imaged using a Nikon eclipse Ti2 microscope using a 4x objective. Subsequently, cancer cells were released from the tissue using trypsin and quantitated by measuring the fluorescence intensity (excitation 577nm, emission 617nm) using SpectraMax iD3 (Molecular Devices). The assay was repeated using tissue collected from 3 patients.

### Bioenergetics assays: Glycolysis and oxidative phosphorylation

The effect of adipocytes and HIF1α on glycolysis and oxidative phosphorylation was determined using the Seahorse XFe analyzer (Agilent, Santa Clara, CA) by measuring extracellular acidification rate (ECAR) and oxygen consumption rate (OCR) ^5, 51^. Briefly, 10,000 cells were plated per well of 96-well Seahorse plates and allowed to attach overnight. Cells were treated with control media or adipocyte-conditioned media for 18hr. Post treatment cells were washed and either replaced with DMEM (D5030, Sigma-Aldrich) supplemented with glutamine (2 mM, Corning, NY) for ECAR analysis and Seahorse DMEM base media (Agilent, Santa Clara, CA), supplemented with sodium pyruvate (1 mM, Corning, NY), glucose (25 mM, Sigma-Aldrich), glutamine (2 mM, Corning, NY) for OCR analysis. The assay media pH was adjusted to 7.4. Cells were incubated in a non-CO2 incubator for 1hr before the assay was started. For ECAR assays, sequential injection of glucose (10 mM, Sigma-Aldrich), Oligomycin (2 μM, Sigma-Aldrich), 2-deoxyglucose (50 mM, Sigma-Aldrich), and for OCR assays, sequential injection with Oligomycin (1 μM, Sigma-Aldrich), FCCP (1 μM, Sigma-Aldrich), Antimycin A (1 μM, SigmaAldrich), Rotenone (1 μM, Sigma-Aldrich) was carried out. Background corrected rate data were plotted to determine changes in ECAR and OCR levels.

### Luciferase activity assay - Oxygen-dependent degradation domain (ODD) – HIF1α stability assay

To quantitatively measure the effect of adipocyte CM HIF1α stability, we transfected SKOV3ip1 and OVCAR5 cells with ODD-luciferase-pcDNA3 construct (a kind gift from William Kaeline, Addgene plasmid # 18965) and generated stable lines ^17^. These stable cells were plated in 6-well plates, allowed to attach for 36hr, and treated with either control media or adipocyte CM in triplicate for 6hr. Cells were subsequently lysed using reporter lysis buffer (Promega), and luciferase activity was measured using an illuminometer (Lumat LB 9507, Berthold technology). Data were normalized to protein levels using a BCA kit (Thermo Fisher Scientific).

### MTT assay

Control shRNA and HIF1α transduced SKOV3ip1 cells were plated in 96 well plates (3000/well) and allowed to attach. The cells were treated with control media (DMEM: F12 media with 1% streptomycin and 1% penicillin) or primary human omental adipocyte-derived conditioned media for 18hr. Subsequently, cells were treated with RSL3 (1 μM), ferrostatin-1 (1μM), and zVAD-fmk (10 μM) and incubated for an additional 6hr. At the end of 24hr MTT assay was carried out as described before ^5^. Relative percent cell viability was calculated based on the values from control media treated group (Csh –Adi control and Hsh – Adi Control).

### Lipid peroxidation assay - Malondialdehyde (MDA)

To determine the intracellular levels of MDA, t**hiobarbituric acid reactive substance assay** was carried out (using TBARS assay kit, Cayman) as described previously **^5^**. Control shRNA and HIF1α transduced SKOV3ip1 cells were plated in 15cm dishes and co-cultured with primary adipocytes (1:5 PCV adipocytes: media) for 18hr. Following which, cell pellets were sonicated and boiled, and assay carried as per manufacturer’s protocol.

### Lipid ROS measurement – C11 Bodipy 581/591

OvCa cells (Control shRNA and HIF1α transduced SKOV3ip1 cells) were plated in black-walled 96 well plates (3000/well) and allowed to attach. The cells were subsequently treated with either control media or adipocyte-derived conditioned media in the presence of (2μM) C11 Bodipy 581/591 dye (Thermo Fisher Scientific) and images were taken every two hours at 10X magnification using IncuCyte S3 live cell imaging system (Sartorius). Cells were continuously imaged for 24hr and the hourly mean total integrated fluorescent intensity measured using the green channel (excitation: 441-481nm, emission – 503-544nm) was plotted for each group.

### Statistical analysis and reproducibility

Graph Pad Prism 7 software was used to calculate statistical significance. Data are presented as mean ± SEM, and a two-tailed student’s t-test was used to determine statistical significance; a p-value less than 0.05 was considered significant. Unless stated, experiments were repeated three times, and one representative experiment was shown. For metabolomic analyses, a mixed-effects one-way ANOVA was carried out. Metabolites with a p-value of < 0.5 were considered significantly changed, and directional changes were depicted as network maps generated using Cytoscape 2.0. Significantly altered metabolites and genes were integrated to obtain a multi-omics dataset. Gene and metabolite pathway over-representation analysis was performed using Integrated Molecular Pathway Level Analysis (IMPaLA)^52^. Enriched pathways were identified based on FDR and adjusted p-values ≤ 0.05. Hierarchical clustering of enriched pathways was subsequently carried out to identify significant networks and visualized using KEGG pathways.

## LIST OF SUPPLEMENTARY MATERIALS

### Figures

Fig. S1. Untargeted metabolomic analysis of adipocytes after co-culture with ovarian cancer cells.

Fig. S2. Joint gene-metabolite analysis of adipocytes co-cultured with cancer cells.

Fig. S3. Seahorse and metabolomics of ovarian cancer cells co-cultured with adipocytes.

Fig. S4. Adipocyte-induced HIF1α regulates glycolysis.

Fig. S5. Adipocyte-induced HIF1α alters the lipidome of ovarian cancer cells.

Fig. S6. Effect of HIF1α knockdown on ovarian cancer cells.

### Tables

Table 1: Differentially expressed genes in both adipocyte and SKOV3i1p OvCa cells after co-culture.

Table 2: Joint gene-metabolite analysis using IMPaLa.

Table 3: Proteomics to determine the effect of HIF1α knockdown on adipocyte induced protein changes.

Table 4: Imaging mass spectrometry.

## AUTHOR CONTRIBUTIONS

A.M and E.L supervised the study, wrote and edited the manuscript. Data acquisition and analysis were carried out by all A.M, D.B, C.C, M.Z, M.T,J.F, D.G, M.R.L, J.W.N, O.F, R.B, and E.L. Adipocyte isolation, co-culture, Seahorse, gene and protein expression analysis (A.M, C.C), Metabolomics (A.M, J.F, M.R.L, D.G), omics data integrations (D.G, A.M), imaging mass spectrometry (F.G, A.M, L.A.M, L.N), [^13^C]-glucose isotope tracing (A.M, D.B, H.S), xenograft studies (A.M).

There are no potential conflicts of interest.

## ACKNOWLEDGMENTS

We thank all patients at The University of Chicago for donating tissue to this study. We thank Drs. Fabian Coscia and Matthias Mann (Max Plank Institute of Biochemistry, Martinsried, Bavaria) for help with proteomic analysis, and J. Andrade at the Center for Research Informatics (University of Chicago) for assistance with microarray analysis. This work was supported by the Ann Sol Schreiber award (372898), Colleen’s dream foundation, and the DOD pilot award (W81XWH2110376) to Abir Mukherjee (A.M), NIH grants (R01CA169604, R35CA264619) awarded to Ernst Lengyel (E.L). D.B is supported by NIH grant F31CA239330. R.J.D. is supported by grants from NIH (R35CA22044901) and by the Howard Hughes Medical Institute. We thank Gail Isenberg for editing the manuscript. Figures 2C, S2E-F, 5B, 6A and F were created using Biorender.

## REFERENCES

1. Lengyel E, Makowski L, DiGiovanni J, Kolonin MG. Cancer as a matter of fat: The crosstalk between adipose tissue and tumors. Trends Cancer 4, 374–384 (2018).

2. Mukherjee A, Bilecz AJ, Lengyel E. The adipocyte microenvironment and cancer. Cancer Metastasis Rev, (2022).

3. Eckert MA, et al. Genomics of ovarian cancer progression reveals diverse metastatic trajectories including intraepithelial metastasis to the fallopian tube. Cancer Discov 6, 1342–1351 (2016).

4. Nieman KM, et al. Adipocytes promote ovarian cancer metastasis and provide energy for rapid tumor growth. Nat Med 17, 1498–1503 (2011).

5. Mukherjee A, et al. Adipocyte-induced FABP4 expression in ovarian cancer cells promotes metastasis and mediates carboplatin resistance. Cancer Res 80, 1748–1761 (2020).

6. Ladanyi A, et al. Adipocyte-induced CD36 expression drives ovarian cancer progression and metastasis. Oncogene 37, 2285–2301 (2018).

7. Lunt SY, Vander Heiden MG. Aerobic glycolysis: meeting the metabolic requirements of cell proliferation. Annu Rev Cell Dev Biol 27, 441–464 (2011).

8. Cluntun AA, Lukey MJ, Cerione RA, Locasale JW. Glutamine metabolism in cancer: Understanding the heterogeneity. Trends Cancer 3, 169–180 (2017).

9. Fagone P, Jackowski S. Membrane phospholipid synthesis and endoplasmic reticulum function. J Lipid Res 50 Suppl, S311–316 (2009).

10. Engelmann B, Wiedmann MK. Cellular phospholipid uptake: flexible paths to coregulate the functions of intracellular lipids. Biochim Biophys Acta 1801, 609–616 (2010).

11. Schug ZT, et al. Acetyl-CoA synthetase 2 promotes acetate utilization and maintains cancer cell growth under metabolic stress. Cancer Cell 27, 57–71 (2015).

12. Buescher JM, et al. A roadmap for interpreting (13)C metabolite labeling patterns from cells. Curr Opin Biotechnol 34, 189–201 (2015).

13. Takeuchi K, Reue K. Biochemistry, physiology, and genetics of GPAT, AGPAT, and lipin enzymes in triglyceride synthesis. Am J Physiol Endocrinol Metab 296, E1195–1209 (2009).

14. Chen HC, et al. Hypoxia Induces a Metabolic Shift and Enhances the Stemness and Expansion of Cochlear Spiral Ganglion Stem/Progenitor Cells. BioMed research international 2015, 359537 (2015).

15. Sumi C, et al. Suppression of mitochondrial oxygen metabolism mediated by the transcription factor HIF-1 alleviates propofol-induced cell toxicity. Sci Rep 8, 8987 (2018).

16. Laurent V, et al. Periprostatic Adipose Tissue Favors Prostate Cancer Cell Invasion in an Obesity-Dependent Manner: Role of Oxidative Stress. Mol Cancer Res 17, 821–835 (2019).

17. Hart PC, et al. Mesothelial Cell HIF1alpha Expression Is Metabolically Downregulated by Metformin to Prevent Oncogenic Tumor-Stromal Crosstalk. Cell reports 29, 4086–4098 e4086 (2019).

18. Safran M, et al. Mouse model for noninvasive imaging of HIF prolyl hydroxylase activity: assessment of an oral agent that stimulates erythropoietin production. Proc Natl Acad Sci U S A 103, 105–110 (2006).

19. Coscia F, et al. Multi-level proteomics identifies CT45 as mediator of chemosensitivity and immunotherapy target in ovarian cancer. Cell 175, 159–170 (2018).

20. Williamson JR, et al. Hyperglycemic pseudohypoxia and diabetic complications. Diabetes 42, 801–813 (1993).

21. Walter M, Liang S, Ghosh S, Hornsby PJ, Li R. Interleukin 6 secreted from adipose stromal cells promotes migration and invasion of breast cancer cells. Oncogene 28, 2745–2755 (2009).

22. Peng G, Liu Y. Hypoxia-inducible factors in cancer stem cells and inflammation. Trends Pharmacol Sci 36, 374–383 (2015).

23. Lengyel E. Ovarian cancer development and metastasis. Am J Pathol 177, 1053–1064 (2010).

24. Yang WS, Stockwell BR. Ferroptosis: Death by Lipid Peroxidation. Trends Cell Biol 26, 165–176 (2016).

25. Yang WS, et al. Regulation of ferroptotic cancer cell death by GPX4. Cell 156, 317–331 (2014).

26. Dixon SJ, et al. Ferroptosis: an iron-dependent form of nonapoptotic cell death. Cell 149, 1060–1072 (2012).

27. Lane AN, Yan J, Fan TW. (13)C Tracer Studies of Metabolism in Mouse Tumor Xenografts. Bio Protoc 5, (2015).

28. Scott AJ, et al. Norharmane Matrix Enhances Detection of Endotoxin by MALDI-MS for Simultaneous Profiling of Pathogen, Host, and Vector Systems. Pathog Dis 74, (2016).

29. Natsume M, et al. Omental adipocytes promote peritoneal metastasis of gastric cancer through the CXCL2-VEGFA axis. Br J Cancer 123, 459–470 (2020).

30. Haverkamp L, Brenkman HJ, Ruurda JP, Ten Kate FJ, van Hillegersberg R. The Oncological Value of Omentectomy in Gastrectomy for Cancer. J Gastrointest Surg 20, 885–890 (2016).

31. Bristow RE, Tomacruz RS, Armstrong DK, Trimble EL, Montz FJ. Survival effect of maximal cytoreductive surgery for advanced ovarian carcinoma during the platinum era: A meta-analysis. J Clin Oncol 20, 1248–1259 (2002).

32. Szachowicz-Petelska B, Dobrzynska I, Skrodzka M, Darewicz B, Figaszewski ZA, Kudelski J. Phospholipid composition and electric charge in healthy and cancerous parts of human kidneys. J Membr Biol 246, 421–425 (2013).

33. Dobrzynska I, Szachowicz-Petelska B, Sulkowski S, Figaszewski Z. Changes in electric charge and phospholipids composition in human colorectal cancer cells. Mol Cell Biochem 276, 113–119 (2005).

34. Piccolis M, et al. Probing the Global Cellular Responses to Lipotoxicity Caused by Saturated Fatty Acids. Mol Cell 74, 32–44 e38 (2019).

35. Listenberger LL, et al. Triglyceride accumulation protects against fatty acid-induced lipotoxicity. Proc Natl Acad Sci U S A 100, 3077–3082 (2003).

36. Petschnigg J, et al. Good fat, essential cellular requirements for triacylglycerol synthesis to maintain membrane homeostasis in yeast. J Biol Chem 284, 30981–30993 (2009).

37. Barrera G. Oxidative stress and lipid peroxidation products in cancer progression and therapy. ISRN Oncol 2012, 137289 (2012).

38. Yang M, et al. Clockophagy is a novel selective autophagy process favoring ferroptosis. Sci Adv 5, eaaw2238 (2019).

39. Zou Y, et al. Plasticity of ether lipids promotes ferroptosis susceptibility and evasion. Nature 585, 603–608 (2020).

40. Mukherjee A. Isolation of Primary Normal and Cancer-Associated Adipocytes from the Omentum (2022).

41. Sellers K, et al. Pyruvate carboxylase is critical for non-small-cell lung cancer proliferation. J Clin Invest 125, 687–698 (2015).

42. Kind T, et al. Fiehnlib: Mass spectral and retention index libraries for metabolomics based on quadrupole and time-of-flight gas chromatography/mass spectrometry. Anal Chem 81, 10038–10048 (2009).

43. Meissen JK, et al. Induced pluripotent stem cells show metabolomic differences to embryonic stem cells in polyunsaturated phosphatidylcholines and primary metabolism. PLoS One 7, e46770 (2012).

44. Olmstead KI, et al. Insulin induces a shift in lipid and primary carbon metabolites in a model of fasting-induced insulin resistance. Metabolomics 13, (2017).

45. Agrawal K, et al. Effects of atopic dermatitis and gender on sebum lipid mediator and fatty acid profiles. Prostaglandins Leukot Essent Fatty Acids 134, 7–16 (2018).

46. Belov ME, et al. Design and Performance of a Novel Interface for Combined Matrix-Assisted Laser Desorption Ionization at Elevated Pressure and Electrospray Ionization with Orbitrap Mass Spectrometry. Anal Chem 89, 7493–7501 (2017).

47. He L, Diedrich J, Chu YY, Yates JR, 3rd. Extracting Accurate Precursor Information for Tandem Mass Spectra by RawConverter. Anal Chem 87, 11361–11367 (2015).

48. Greco F, et al. Mass Spectrometry Imaging as a Tool to Investigate Region Specific Lipid Alterations in Symptomatic Human Carotid Atherosclerotic Plaques. Metabolites 11, (2021).

49. Smith CA, et al. METLIN: a metabolite mass spectral database. Ther Drug Monit 27, 747–751 (2005).

50. Liebisch G, et al. Shorthand notation for lipid structures derived from mass spectrometry. J Lipid Res 54, 1523–1530 (2013).

51. Curtis M, et al. Fibroblasts mobilize tumor cell glycogen to promote proliferation and metastasis. Cell Metab 29, 141–155 e149 (2019).

52. Kamburov A, Cavill R, Ebbels TM, Herwig R, Keun HC. Integrated pathway-level analysis of transcriptomics and metabolomics data with IMPaLA. Bioinformatics 27, 2917–2918 (2011).

